# High extracellular lactate increases reductive carboxylation in breast tissue cell lines grown under normoxic conditions

**DOI:** 10.1101/558296

**Authors:** Arthur Nathan Brodsky, Daniel C. Odenwelder, Sarah W. Harcum

## Abstract

In cancer tumors, lactate accumulation was initially attributed to high glucose consumption associated with the Warburg Effect. Now it is evident that lactate can also serve as an energy source in cancer cell metabolism. Additionally, lactate has been shown to promote metastasis, generate gene expression patterns in cancer cells consistent with “cancer stem cell” phenotypes, and result in treatment resistant tumors. Therefore, the goal of this work was to quantify the impact of lactate on metabolism in three breast cell lines (one normal and two breast cancer cell lines – MCF 10A, MCF7, and MDA-MB-231), in order to better understand the role lactate may have in different disease cell types. Parallel labeling metabolic flux analysis (^13^C-MFA) was used to quantify the intracellular fluxes under normal and high extracellular lactate culture conditions. Additionally, high extracellular lactate cultures were labelled in parallel with [U-^13^C] lactate, which provided qualitative information regarding the lactate uptake and metabolism. The ^13^C-MFA model, which incorporated the measured extracellular fluxes and the parallel labeling mass isotopomer distributions (MIDs) for five glycolysis, four tricarboxylic acid cycle (TCA), and three intracellular amino acid metabolites, predicted lower glycolysis fluxes in the high lactate cultures. All three cell lines experienced increased reductive carboxylation of glutamine to citrate in the TCA cycle as a result of high extracellular lactate. Increased reductive carboxylation previously has been observed under hypoxia and other mitochondrial stresses, whereas these cultures were grown aerobically. In addition, this is the first study to investigate the intracellular metabolic responses of different stages of breast cancer progression to high lactate exposure. These results provide insight into the role lactate accumulation has on metabolic reaction distributions in the different disease cell types while the cells are still proliferating in lactate concentrations that do not significantly decrease exponential growth rates.

## Introduction

Since the 1920s, many types of cancers have been shown to rely heavily on glycolysis and lactate fermentation to produce energy rather than the more energy efficient complete oxidation of glucose in the mitochondria, even in the presence of sufficient oxygen. This metabolic state is called the Warburg Effect [1-4]. In addition, lactate can be utilized by cancer cells in the presence of glucose, a process known as the Reverse Warburg Effect [5-9]. Not only does this capability to use lactate provide cancer cells a metabolic advantage *in vivo*, it seems to favor cancer progression. For example, when lactate was injected into mice with xenografts of the human breast cancer cell line MDA-MB-231, metastasis increased ten-fold [10]. When the human breast cancer cell line MCF7 was exposed to lactate *in vitro*, genes associated with “stemness” were upregulated and gene expression patterns consistent with the “cancer stem cell” phenotype were observed [7]. In several other types of cancers, intratumoral lactate levels – which can rise to as high as 40 mM – correlated with treatment resistance as well as poor patient prognosis [5]. Further, it has been shown in xenotransplants and mouse cancer models that inhibiting the ability of cancer cells to utilize lactate can force the cells to become glycolytic and retard tumor growth through glucose starvation, while rendering the remaining cells more susceptible to radiation treatments [9]. Since lactate accumulation and its subsequent utilization by surrounding cancer cells appears to negatively affect cancer patient outcomes, deciphering the role of lactate at the metabolic level within central carbon metabolism is crucial.

Metabolic flux analysis (MFA) is a computational tool that is used to quantify the intracellular metabolic fluxes of individual metabolic pathways [11, 12]. MFA can be used to compare changes in metabolic activity due to particular factors [13, 14]. Classical MFA uses extracellular uptake and secretion rates of nutrients and waste products, and a discrete metabolic network model of the relevant metabolic reactions [15, 16]. The use of stable isotopic tracers, typically ^13^C-labeled nutrients, and the resulting mass isotopomer distribution (MID) data increases the resolution of individual fluxes within the defined metabolic reaction network [17-19]. Several mammalian systems have been characterized including Chinese hamster ovary (CHO) cells at both stationary and exponential growth phases [20-22], MDA-MB-231 cells under various nutrient conditions [23], and several other cancer cell lines under hypoxia [24, 25]. There are several software tools available to assist researchers with resolving the intracellular fluxes including Metran, OpenFLUX, 13CFlux2, INCA and FiatFlux [22, 26-29]. All of these software tools rely upon regression analysis to solve the system of linear equations specified by the metabolic network.

The aim of this study was to determine the role of extracellular lactate on the metabolism of three different proliferating breast cell lines (one non-tumorigenic epithelial breast cell line and two breast cancer cell lines). Each cell line was grown in a low glucose control (e.g., normal or typical laboratory) media or in a high-lactate media, where sodium lactate was added to the control medium to make the high-lactate medium. The high-lactate concentration was selected for each cell line such that equivalent exponential growth rates for the control and high-lactate cultures were maintained. These equivalent exponential growth rates were important for the modeling assumptions for two reasons: 1) the quasi-steady state assumption was met equally well by both conditions, and 2) it allowed for normalizations of fluxes between the conditions for the same cell line based on biomass generation. Parallel labeling experiments were conducted with [1,2-^13^C] glucose, [U-^13^C] L-glutamine, and [U-^13^C] sodium lactate, where the control cultures were only labeled with [1,2-^13^C] glucose and [U-^13^C] L-glutamine. The lactate labeling data for the high-lactate cultures provided uptake and direct intermediate tricarboxylic acid (TCA) cycle labeling information. The intracellular metabolic fluxes were predicted for each cell line and each condition from the glucose and glutamine labeling data.

## Materials and methods

### Cell lines and media formulations

MCF 10A (ATCC^®^ CRL-10317^TM^), MCF7 (ATCC^®^ HTB-22^TM^) and MDA-MB-231 (ATCC^®^ HTB-26^TM^) cells were from the American Tissue Culture Collection (ATCC). MCF 10A cells are a non-tumorigenic breast cell line, MCF7 cells are a tumorigenic, luminal breast cancer cell line, and MDA-MB-231 cells are a metastatic, basal breast cancer cell line [30]. Dulbecco’s modified Eagle medium (DMEM) without glucose, glutamine, sodium pyruvate, and phenol red (Life Technologies) was used as the growth media. The growth media was supplemented to contain 5 mM glucose (Thermo Fisher), 3 mM glutamine (Life Technologies), 10% dialyzed fetal bovine serum (dFBS, Life Technologies), 100 U/mL penicillin, and 100 μg/mL streptomycin (100X Penicillin-Streptomycin solution, Life Technologies). High-lactate cultures were supplemented with 10 mM sodium L-lactate (Sigma) for the MCF 10A cultures and 20 mM for the MCF7 and MDA-MB-231 cultures. The isotopic tracers were [1,2-^13^C] glucose, [U-^13^C] L-glutamine, and [U-^13^C] sodium lactate, all purchased from Sigma-Aldrich.

### Cell growth and parallel labeling experiments

All cell lines were seeded at 2 × 10^4^ cells/cm^2^ into either 6-well plates or T-25 flasks in the control growth media. Cells were cultured at 37°C in a 5% CO_2_ humidified incubator. After 24-h, the media for the MDA-MB-231 cultures was exchanged and the experimental conditions were introduced. The media exchange time is used to set time 0 for all cell lines. For MCF 10A and MCF7 cell lines the lag phase was longer, so the media was replenished at 24-h post seeding. The media was exchanged 48-h post seeding for MCF10A and MCF 7 to introduce the experimental conditions. For clarity, the full experimental setup is shown in Fig 1. Cell numbers and extracellular metabolite concentrations were measured as shown in Fig 1, where all times are relative to the isotope media exchange. Cell numbers and glucose and lactate concentrations were obtained from the six-well plates, while the amino acid concentrations were obtained from the T-25 flasks. Cells for intracellular metabolite analysis were obtained from the T-25 flasks 24-h after the media exchange, since 24-h has previously been shown to provide sufficient amount of time for isotopic steady state to be achieved for intracellular glycolytic metabolites in CHO cells [21]. Parallel cultures of glucose (95% molar enriched [1,2-^13^C] glucose) and glutamine tracers (95% molar enriched [U-^13^C] L-glutamine) were performed for the control cultures, whereas the high-lactate cultures also had parallel replicates for the lactate tracer (50% molar enriched [U-^13^C] sodium L-lactate). Six-well plates (Nunc) were used to obtain cell numbers and glucose and lactate concentrations with six replicates for each time point. T-25 flasks were used to obtain extracellular amino acid concentrations and intracellular MID measurements in triplicate.

**Fig 1.**
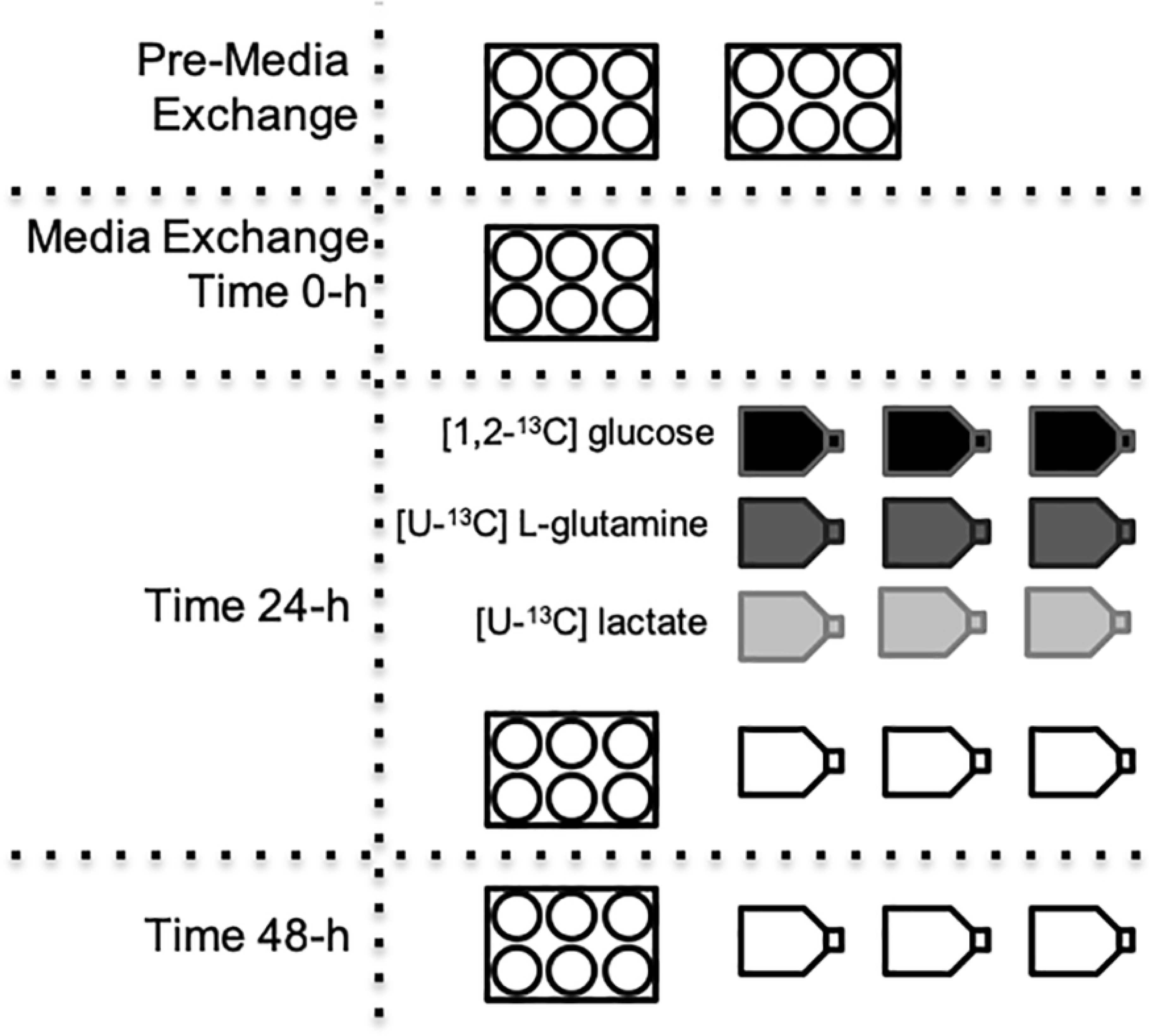
Experimental setup for the parallel labeling experiments for each of the three breast cell lines. The four or five six-well plates and 15 T-25 flasks were seeded simultaneously for each cell line examined. The times shown indicate when the plates or flasks were harvested for analysis relative to the isotope media exchange. Isotopic labeling is graphically shown by shading (white – no isotope, black – glucose, dark grey – glutamine, and light grey – lactate). The six-well plates did not contain isotopically-labeled media.

### Cell numbers, and glucose and lactate concentrations

Cell numbers were obtained using the Scepter 2.0 Handheld Automated Cell Counter (Millipore). Glucose and lactate concentrations were measured using a YSI 2700 Bioanalyzer.

### Preparation of samples for gas chromatography-mass spectrometry analysis

Intracellular and extracellular metabolites and specific metabolite fragments used for identification were extracted and derivatized according to the protocol outlined by Ahn and Antoniewicz [22], with a few minor changes. Samples were incubated and derivatized for 60 min, and the final sample volumes were increased to 1 mL after derivatization by adding additional pyridine. The injection volumes were 3 μL and samples were injected in splitless mode. Amino acid standards were used to calculate amino acid concentrations in the [U–^13^C] algal amino acid solution, which were then used to quantify the amino acid concentrations in the extracellular medium.

### Extracellular amino acid concentrations and intracellular MID measurements

GC-MS analysis was performed using a Hewlett Packard 7683 GC equipped with an HP-5 (30 m × 0.32 mm i.d. × 0.25 μm; J&W Scientific) capillary column, interfaced with a Hewlett Packard 5973 MS operating under ionization by electron impact as 2000 eV and 200°C ion source temperature. The injection port and interface temperatures were both 250°C while helium flow was maintained at 1 mL/min. Mass spectra were recorded in full scan mode for amino acid quantification and in single ion mode (SIM) for MIDs as well as internal standards and standardization curves. MIDs were obtained by integration of single ion chromatograms and corrected for natural isotope abundances using the Metran software [31, 32].

### Determination of biomass specific consumption and production rates

Specific consumption and production rates for nutrients and waste metabolites in the extracellular medium were determined based on cell growth rate and nutrient time profiles, where the times are relative to media exchange and not to seeding [33]. For each sample calculation, statistical analysis of the amino acid concentration data was conducted using JMP 10.0.0 (SAS Institute, Inc.). The generalized linear model was used where cell line, condition, and time were examined as effectors of the amino acid concentration (p ≤ 0.05). All amino acid concentrations were determined to be significantly affected by time (p ≤ 0.05). Several amino acid concentrations were not significantly affected by cell line, condition, or both (p > 0.05). The spontaneous glutamine degradation rate was accounted for in the glutamine flux calculation, with a degradation rate of 0.0019 h^-1^ [33].

### Metabolic network model

To model central carbon metabolism for the three breast cell lines, a general mammalian cell model was used. This generalized mammalian cell model was adapted from the previously developed framework by Ahn and Antoniewicz (2013). Major reactions for glycolysis, the pentose phosphate pathway (PPP), the TCA cycle, amino acid metabolism, lactate metabolism, and fatty acid metabolism were included in the mammalian cell model. Carbon flux to cellular biomass was broken down into two separate compartments, such that the potentially higher lipid content of the breast cells could be modeled. The Biomass pool includes proteins, nucleotides, and carbohydrates and the Lipid Biomass pool includes lipids and phospholipid [21, 24, 34]. Several studies have shown that both total lipid weight fraction and lipid content vary between breast tissue and tumor tissue layers [35-37]. The initial flux to the Lipid Biomass was constrained to 10% of the sum of the total cell mass [33]. In order to reduce the complexity of the model to align with the number and detail of measurements taken, only citrate and acetyl-CoA were modeled with both cytosolic and mitochondrial compartments [18, 21, 24]. A lactate sink species was included to account for the dilution of labeled lactate by metabolism of unlabeled glucose [38]. Carbon dioxide was treated as an unbalanced metabolite and was not measured. Oxygen uptake was excluded from the model and was also not measured. Cofactor balances, such as NADH and NADPH were not included in the model, as different isozymes have varying cofactor requirements and the inclusion of these assumptions can skew subsequent analysis [22]. The breast cell metabolic flux model has 38 reactions. The full breast cell metabolic flux model with atom transitions is included in Table D in S1 Appendix.

### Metabolic flux analysis

^13^C-Metabolic flux analysis was performed using the software package Metran [32], which utilizes the elementary metabolite unit (EMU) framework [39]. Metabolic fluxes were estimated using experimentally measured-values for extracellular consumption and production rates and from MIDs obtained for intracellular metabolites. In Metran, the intracellular and extracellular metabolic flux predictions were based on quantities that minimized the variance-weighted sum of squared residuals (SSRes) between the measured values input into the model and the simulated values. Metran can process MID data from parallel labeling experiments to predict the metabolic fluxes, a capability that has been validated using both *E. coli* and CHO cells experimental data [21, 38]. In this study, random initial fluxes were used, and the MID error was calculated from the biological replicates. Additionally, a minimum MID error threshold of 0.6 mol% was applied if the biological error was less than 0.6 mol%; this falls within the standard error range that has been used in previous MFA studies [18, 21]. Most of the standard errors observed in this study for the biological replicates were higher than the 0.6 mol% machine error used previously when replicates were not available [21]. The MIDs metabolites labeled by [1,2-^13^C] glucose included in the MFA simulations were 3-phosphoglycerate (3PG), dihydroxyacetone phosphate (DHAP), pyruvate, lactate, and alanine. The MIDs metabolites labeled by [U-^13^C] glutamine included in the MFA simulations were succinate, malate, α-ketoglutarate (AKG), glutamate, citrate, glutamine, and pyruvate. The MIDs of metabolites labeled by [U-^13^C] lactate were not included in the MFA simulations.

The extracellular flux for each metabolite was calculated as described in Meadows et al. (2008) [33], and adapted to the media exchange time as:

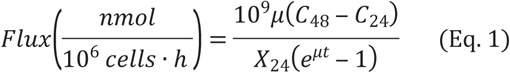

where, *μ* is the growth rate (ℎ^‒1^). *C*_48_ and *C*_24_ are the individual metabolite concentrations (mM) at 48 and 24 hours after the media exchange, respectively. *X*_24_ is the cell concentration (cells/mL) at 24 hours after the media exchange. In Meadows et al. (2008) [33], only the term X_0_ was used in the flux equation instead of X_24_, as their flux calculation included the initial time, time 0. In the current study, the flux calculation only includes the time points 24-h and 48-h, thus the cell concentration uses the 24-h time point as the reference point for the cell concentration. The value for the term *t* is 24 hours for all of the current flux calculations.

### Statistical Analysis

Statistical analysis was performed using the software JMP pro 10 (SAS Institute, Cary, NC). The generalized linear model (GLM) procedure (p ≤ 0.05) and least squares method (LS mean) with Tukey HSD (honestly significant difference) were used to determine if growth rates, cell numbers 24-h after the medium exchange, glucose, lactate, and amino acid concentrations were affected by the cell line and/or condition. To estimate the standard deviation for each metabolite flux, Monte Carlo simulations were conducted using 1,000,000 iterations of the flux equation, where the standard deviation for each input *μ*, (*C*_48_ ‒ *C*_24_), and *X*_24_ was applied. For Metran, metabolic flux simulations were determined to have converged when a global solution was achieved that satisfied the accepted SSRes criteria, unless otherwise specified. This was determined from analyzing the simulated fit results via a chi-square statistical test to measure goodness-of-fit [40, 41]. After convergence, 95% confidence intervals were generated for all parameters based on the SSRes parameter [40].

## Results

### Cell growth

To determine the effects of high extracellular lactate on breast cancer metabolism, three human breast cell lines, MCF 10A, MCF7, and MDA-MB-231, were grown under both control and high-lactate conditions. MCF 10A is a non-tumorigenic epithelial cell line, and MCF7 and MDA-MB-231 are two different stages of breast cancer (tumorigenic, luminal and metastatic, basal respectively). Sodium lactate was used for the high-lactate conditions to reduce the initial media pH shift and prevent shocking the cells due to the lactate addition. For MCF7 and MDA-MB-231, the high-lactate culture media consisted of the control media supplemented with 20 mM sodium lactate. For MCF 10A, the high-lactate culture media consisted of the control media supplemented with 10 mM sodium lactate. MCF 10A cells were unable to grow under the 20 mM lactate stress; however, MCF 10A could grow at equal growth rates to the control culture in 10 mM lactate-supplemented media. Both 10 mM and 20 mM lactate represent normal physiological concentration ranges [5-7]. Regrettably, the MCF 10A high-lactate cultures had to be cultured at a lower lactate concentration than the MCF7 and MDA-MB-231 high-lactate cultures, which prevented direct comparisons across all three cell lines. Yet, maintaining equal growth rates between the control and high-lactate conditions within each cell line was central to the experimental design. This allowed for any observed metabolic changes to be attributed to media condition alone and not to a shift in growth rate.

The control media was composed of Dulbecco’s modified Eagle medium (DMEM) with a lower initial glucose concentration (5 mM) than standard DMEM (25 mM), in order to be more representative of physiological glucose concentrations, where 5 mM is equal to 0.90 g/L or 90 mg/dL. To minimize the interference of other carbon sources on the labeling of the intracellular metabolites, other potential carbon sources in the media were reduced or eliminated. For example, sodium pyruvate was eliminated from the standard DMEM formulation, and dialyzed fetal bovine serum (FBS) was used to eliminate glucose and glutamine carryover and reduce the unquantified amino acids from FBS. These modifications to the media allowed for the labeling studies to be conducted with 95% labeled glucose and glutamine.

To characterize cell growth, cell counts were taken every 24-h; growth profiles for the three cell lines are shown in Fig 2. All three cell lines exhibited significant reproducible lag phases when passaged in the control media, likely due to the lower than normal glucose concentration, despite previously being adapted to this media formulation for over three passages. The lag phases were 48-h for MCF 10A and MCF7 and 24-h for MDA-MB-231. After the lag phases, the culture media was exchanged to either fresh control media or the high-lactate media, as indicated at time 0 in Fig 2. For the labeling studies, ^13^C isotopes replaced the unlabeled glucose, glutamine, or lactate in the parallel cultures, at the same concentrations. After the lag phase and media exchange, cells from all culture conditions maintained exponential growth for 48-h, confirming that the high-lactate media did not significantly affect the growth rates of any cell line (p > 0.05). The exponential growth rates were 0.017 h^-1^, 0.018 h^-1^, and 0.021 h^-1^ for the MCF 10A, MCF7, and MDA-MB-231 cell lines, respectively. Previously, 24-h was shown to be sufficient amount of time for isotopic steady-state to be reached for glycolysis and PPP metabolites from glucose labeling, while TCA metabolites also approach isotopic steady state from glutamine labeling within 24-h for mammalian cells [18, 21]. Therefore, in these studies, samples for intracellular MID analysis were taken 24-h after ^13^C-labeled media addition, indicated by the black boxes. The experiments were designed such that all cultures were in the mid-exponential phase at the time of harvest for intracellular MID analysis (Fig 2); this represents a pseudo-steady state, a condition consistent with the MFA assumptions [11].

**Fig 2.**
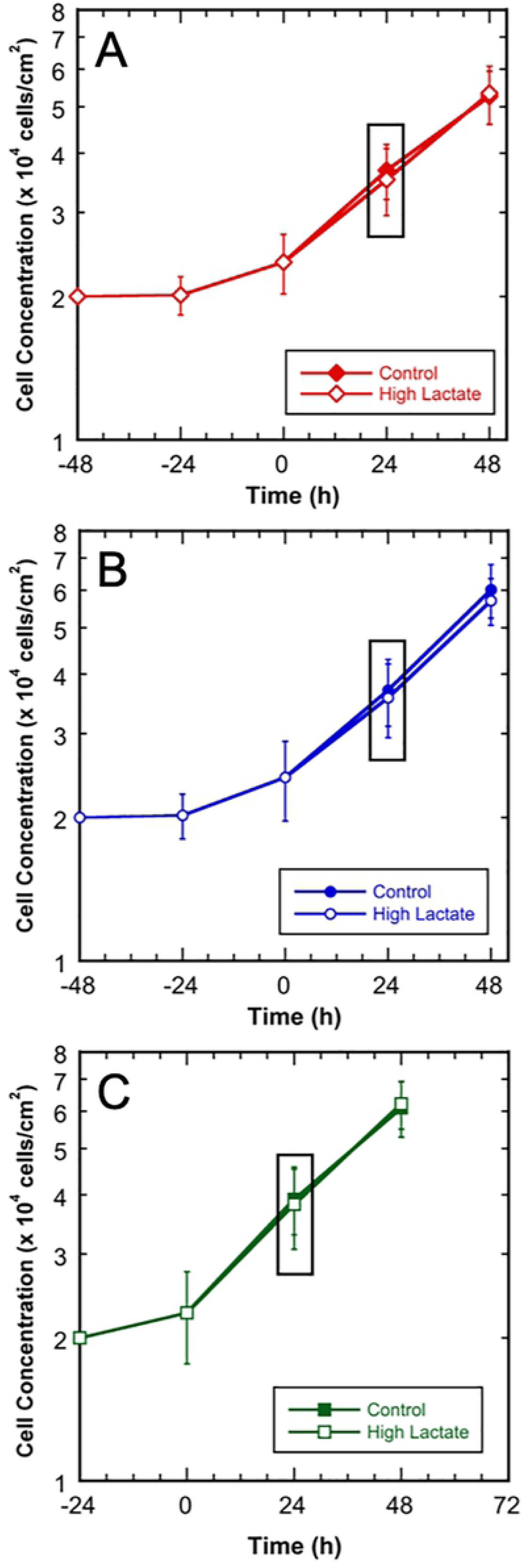
Growth profiles for MCF 10A, MCF7, and MDA-MB-231 cultures in control and high-lactate media. The isotope media exchange occurred at 0-h and samples for MID analysis were taken at 24-h. 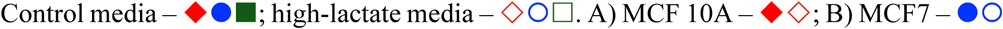 and 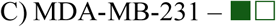. Error bars represent standard deviations.

### Glucose, lactate, and amino acid metabolism

The control and high-lactate conditions exhibited equal growth rates within each cell line; however, high extracellular lactate resulted in reduced glucose utilization and reduced lactate accumulation for each cell line (p ≤ 0.05). Fig 3 shows the glucose and lactate concentration time profiles for each cell line and condition. The average glucose and lactate concentrations for all the 24-h and 48-h samples are listed with the standard deviations in Tables A-C in S1 Appendix. Glucose and lactate fluxes were calculated using concentrations measured at 24-h and 48-h with the cell number at 24-h and the growth rates as per Eq.1 and are listed with standard deviations in Table 1. The standard deviations for the glucose and lactate fluxes were determine using Monte Carlo simulations due to sample independence [42]. The three control cultures exhibited higher glycolytic efficiencies compared to the high lactate cultures. Specifically, the glycolytic efficiencies for the control conditions were 1.7, 1.5, and 1.8 moles lactate produced per mole glucose consumed, respectively, for the MCF 10A, MCF7, and MDA-MB-231 cell lines. The glycolytic efficiencies for the high-lactate conditions were 1.4, 0.8 and 1.3 moles lactate produced per mole glucose consumed, respectively, for MCF 10A, MCF7, and MDA-MB-231 cell lines. In comparison, the theoretical maximum glycolytic efficiency is 2.0 moles lactate produced per mole glucose consumed [30].

**Table 1.** Measured extracellular fluxes for glucose, lactate, and amino acids for the MCF 10A MCF7, and MDA-MB-23 cell lines in the control and high-lactate media. Standard deviations were determined using Monte Carlo simulations of the flux equation due to sample independence [30]. Negative values represent consumption rates and positive values represent production rates.

| Flux (nmol/10<sup>6</sup> cells·h) |  |  |  |  |  |  |  |  |  |  |  |  |
| --- | --- | --- | --- | --- | --- | --- | --- | --- | --- | --- | --- | --- |
| Cell line | MCF 10A |  |  |  | MCF7 |  |  |  | MDA-MB-231 |  |  |  |
| Condition | Control |  | High-Lactate |  | Control |  | High-Lactate |  | Control |  | High-Lactate |  |
| Metabolite | Flux | SD | Flux | SD | Flux | SD | Flux | SD | Flux | SD | Flux | SD |
| Glucose (N=6) | -222 | 51 | -182 | 48 | -208 | 44 | -158 | 42 | -337 | 65 | -267 | 67 |
| Lactate (N=6) | 379 | 56 | 254 | 55 | 318 | 54 | 131 | 32 | 603 | 108 | 342 | 87 |
| Glutamine (N=3) | -48 | 18 | -55 | 19 | -53 | 19 | -63 | 18 | -44 | 21 | -55 | 21 |
| **Amino Acid (N=3)** |  |  |  |  |  |  |  |  |  |  |  |  |
| Alanine | 11 | 1.5 | 11 | 1.9 | 14 | 2.3 | 12 | 2.3 | 10 | 1.8 | 8.2 | 2.0 |
| Aspartate | 4.9 | 0.8 | 3.7 | 0.7 | 6.5 | 1.3 | 5.4 | 1.1 | 3.5 | 0.7 | 4.5 | 1.1 |
| Glutamate | 12 | 1.9 | 10 | 1.8 | 11 | 2.0 | 12 | 2.3 | 8.5 | 1.6 | 10 | 2.5 |
| Glycine | 6.2 | 1.6 | 5.7 | 3.1 | 9.6 | 3.0 | 9.4 | 3.2 | 7.8 | 2.1 | 7.0 | 2.6 |
| Isoleucine | -8.3 | 2.3 | -8.4 | 2.0 | -9.5 | 2.0 | -9.2 | 2.3 | -4.7 | 1.6 | -8.4 | 2.4 |
| Leucine | -6.6 | 2.1 | -7.5 | 2.7 | -8.3 | 2.7 | -7.5 | 2.2 | -4.7 | 2.1 | -8.8 | 2.4 |
| Methionine | -2.6 | 1.0 | -1.5 | 0.8 | -3.1 | 1.1 | -2.2 | 0.8 | -1.4 | 0.5 | -2.2 | 1.0 |
| Phenylalanine | -4.7 | 1.9 | -4.4 | 1.5 | -3.8 | 1.3 | -3.8 | 1.5 | -3.4 | 0.9 | -4.0 | 2.0 |
| Proline | 1.5 | 0.3 | 1.3 | 0.3 | 2.8 | 0.5 | 2.3 | 0.5 | 0.8 | 0.2 | 1.3 | 0.3 |
| Serine | -19 | 2.9 | -20 | 4.0 | -20 | 4.0 | -23 | 5.2 | -16 | 3.4 | -20 | 5.0 |
| Threonine | -1.9 | 1.1 | -1.9 | 0.9 | -1.2 | 0.5 | -1.2 | 0.6 | -1.2 | 0.6 | -1.6 | 0.9 |
| Tyrosine | -3.6 | 1.5 | -3.7 | 1.9 | -4.0 | 1.2 | -3.8 | 1.5 | -3.5 | 1.6 | -3.8 | 1.8 |
| Valine | -5.8 | 1.7 | -6.2 | 1.6 | -9.4 | 2.0 | -10 | 3.3 | -6.3 | 1.7 | -9.1 | 2.3 |

**Fig 3.**
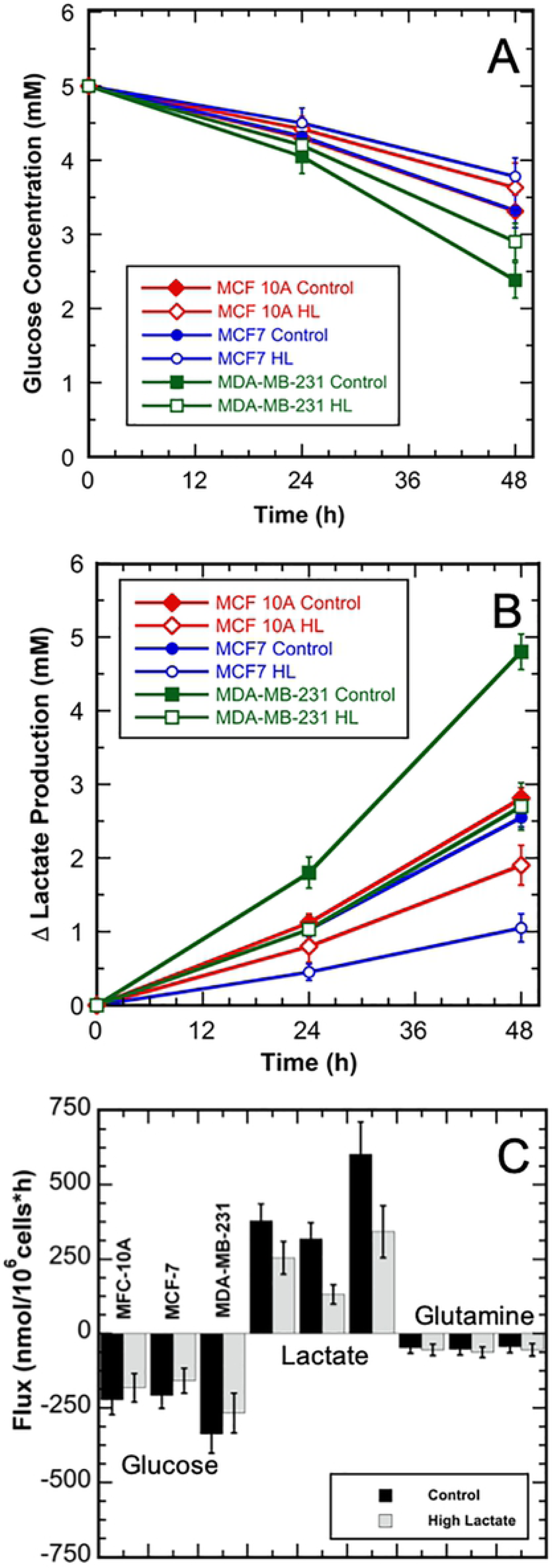
Glucose and lactate concentration and glucose, lactate, and glutamine flux profiles for the three breast cell line cultures in the control and high-lactate media shown relative to the time of the media exchange. A) Glucose concentrations; B) Change in lactate concentrations. 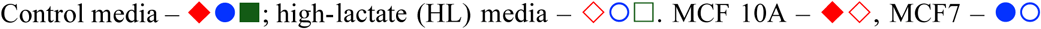 and 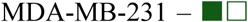. C) Glucose, lactate, and glutamine fluxes. 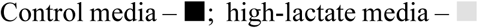. All error bars represent standard deviations.

Amino acid metabolism plays an important role in protein and nucleotide synthesis, and anaplerosis for mammalian cells. Extracellular amino acid concentrations were measure at 24-h and 48-h after the media exchange to quantify amino acid fluxes. Prior to calculating the amino acid fluxes, statistical analysis was conducted for the amino acid concentrations with respect to culture condition, cell line, and time. All the amino acid concentrations were statistical different with respect to time (p ≤ 0.05), such that fluxes could be calculated; however, many of the amino acid concentrations were not statistically different between the cell lines or culture condition (p > 0.05). For example, glutamine (Fig 3C), methionine, phenylalanine, and tyrosine concentrations were not significantly different between cell lines or between the culture conditions (p > 0.05). Whereas, aspartate, glutamate, glycine, and threonine concentrations were significantly different between cell lines (p ≤ 0.05), but were not significantly different within the same cell line between the control and high-lactate conditions (p > 0.05). In addition, leucine concentrations were not significantly different between cell lines (p > 0.05), but were significantly different between the control and high-lactate conditions (p ≤ 0.05). Table 1 lists the calculated amino acid fluxes with the standard deviations determined using Monte Carlo simulations due to sample independence [42]. The average amino acid concentrations for all the 24-h and 48-h samples with the standard deviations are shown in Tables A-C in S1 Appendix.

### Lactate tracer uptake

To further understand the role of lactate as a metabolic substrate, cells were grown with ^13^C-labeled lactate for the high extracellular lactate conditions. This allowed for intracellular metabolite labeling due to lactate uptake and metabolism. Labeled intracellular metabolites measured included: lactate, pyruvate, alanine, citrate, AKG, glutamate, succinate, fumarate, malate, and aspartate (Fig 4A). The intracellular lactate MIDs from [U-^13^C] lactate labeling of the MCF 10A, MCF7, and MDA-MB-231 cultures exposed to 10 mM, 20 mM, and 20 mM extracellular lactate, respectively, are shown in Fig 4B. MCF7 cells had the highest intracellular lactate isotopomer labeling. The TCA metabolites had appreciable levels of M1 labeling in all cell lines. These results indicate that extracellular lactate was consumed by the cells even in the presence of glucose. For completeness, Fig A in S1 Appendix illustrates the intermediate TCA metabolite labeling from [U-^13^C] lactate, as the primary carbon source, as well as the TCA carbon labeling from unlabeled pyruvate and dilution of the labeled metabolite pools. All raw uncorrected lactate-derived MIDs for the three cell lines and for both conditions are provided in Tables E and F in S1 Appendix.

**Fig 4.**
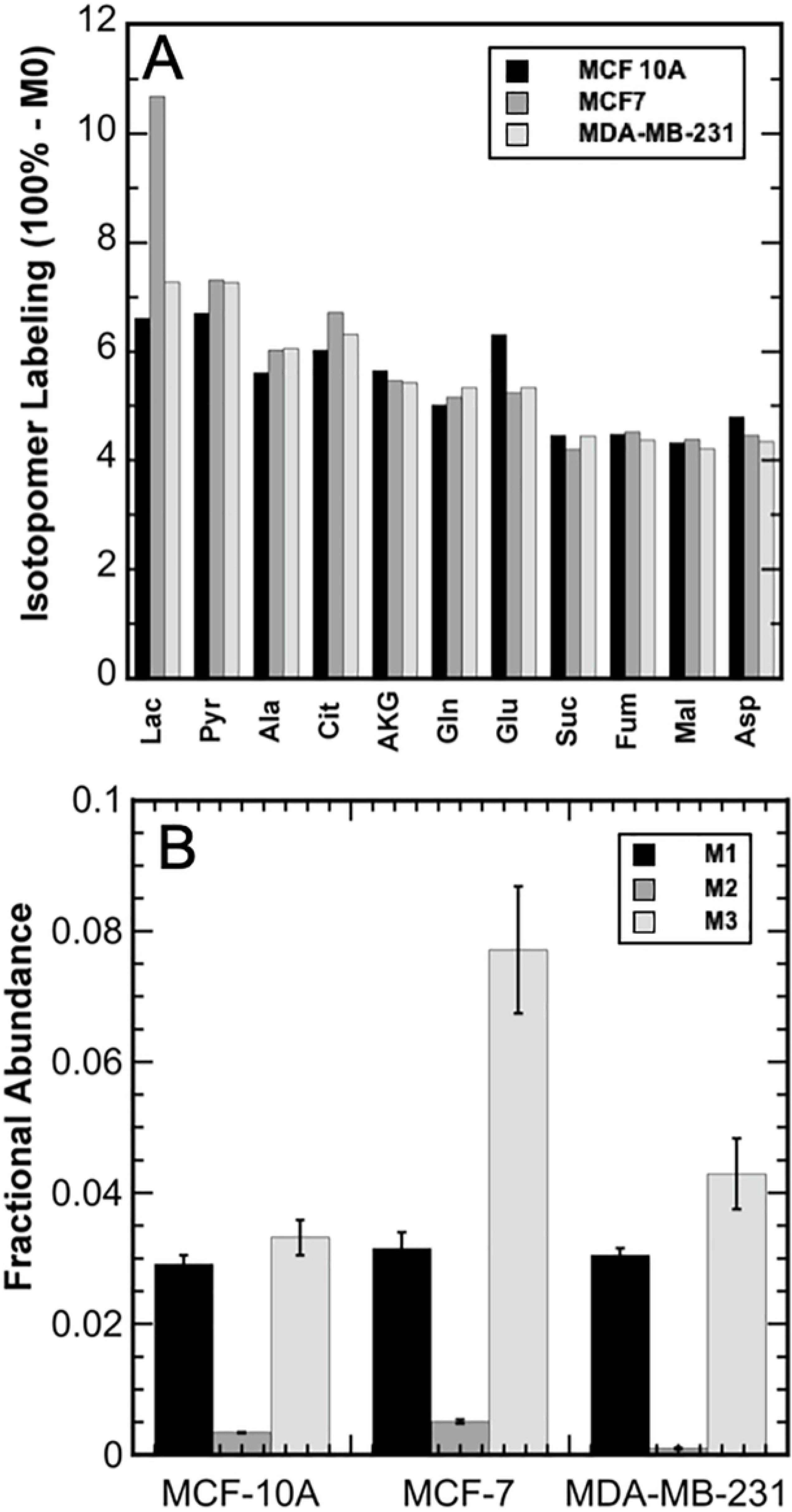
Uptake of U-^13^C lactate by the three breast cell lines. A) Percent carbon labeling for intracellular metabolites due to extracellular [U-^13^C] lactate in the media. 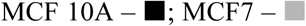 and 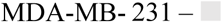. B) Lactate intracellular mass isotopomer distributions (MIDs) due to extracellular [U-^13^C] lactate. 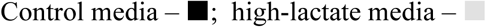. Error bars represent standard deviations.

### Intracellular labeling from [1,2-^13^C] glucose and [U-^13^C] glutamine

[1,2-^13^C] glucose and [U-^13^C] glutamine isotopes were used for the parallel labeling studies to characterize the relative metabolic contribution of each substrate and estimate intracellular metabolic fluxes for glycolysis, the PPP, and the TCA cycle. These isotopes have previously been shown to be suitable tracers for characterizing fluxes for mammalian cells through glycolysis and the PPP ([1,2-^13^C] glucose) and the TCA cycle ([U-^13^C] glutamine) pathways [18]. The percent isotopomer labeling (100% - M0) was calculated for each metabolite using MIDs corrected for natural abundance [21, 31]. Using [1,2-^13^C] glucose, the percent isotope labeling for DHAP, 3PG, pyruvate, alanine, and lactate were 40% or higher for all three cell lines. No significant differences were observed for MIDs from glycolytic metabolites (DHAP, 3PG, pyruvate, alanine, and lactate) between the three cell lines and between the control and high-lactate conditions. These results reflect the consistently high flux through glycolysis for each of the three cell lines and the equivalent growth rates between conditions within a given cell line. These results also confirm that the 24-h labeling time was sufficient to reach isotopic steady state for [1,2-^13^C] glucose. The intracellular MID profiles for all three cell lines are included in Fig B in S1 Appendix. All raw uncorrected MIDs for each cell line and condition are provided in Tables E and F in S1 Appendix. Additionally, all the measured MIDs, corrected for natural isotope abundance, are shown in Figs B and C in S1 Appendix.

For the three cell lines and both conditions, glutamine, AKG, and glutamate were highly labeled from [U-^13^C] glutamine. Using [U-^13^C] glutamine, the observed isotopomer labeling for intracellular glutamine, AKG and glutamate was over 70%. Fig 5 shows the control and high lactate intracellular MID profiles for MDA-MB-231 from [U-^13^C] glutamine, as an example, for the TCA metabolites. Pyruvate and lactate were minimally enriched from the glutamine isotope across the cell lines and both conditions, where Fig 5 shows the MDA-MB-231 data. All the MID profiles for the three cells lines for both conditions are shown in Fig C in S1 Appendix. For MCF 10A and MDA-MB-231, the high-lactate condition resulted in a higher percent isotope labeling of pyruvate and lactate as compared to the control condition. For MCF7, the trend was reversed – the percent isotope labeling of pyruvate and lactate were lower in high-lactate condition compared to the control condition.

**Fig 5.**
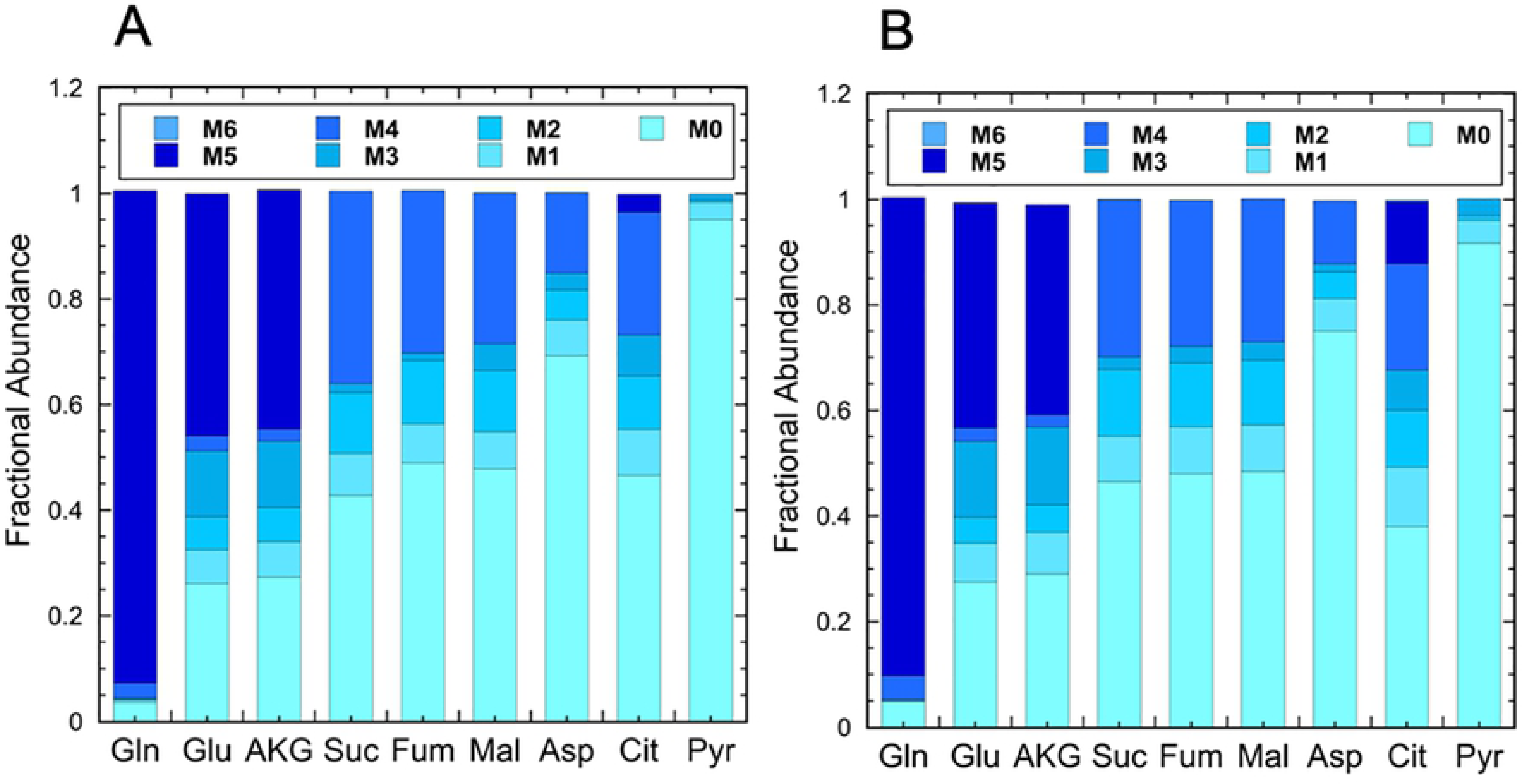
Mass isotope distributions (MIDs) for the TCA metabolites from the [U-^13^C] glutamine labeling of the MDA-MB-231 cells cultured in the control and high-lactate media. A) Control media; B) High-lactate media.

The TCA metabolites succinate, fumarate, malate, and citrate were all significantly labeled by the [U-^13^C] glutamine tracer. The MID profiles for these metabolites were similar for all three cell lines; however, citrate MIDs were significantly different between the control and high-lactate cultures within a cell line, indicating the effect of lactate was significant. More specifically, a significant increase in M5 citrate labeling was observed in the high-lactate conditions for all three cell lines (Fig 6A). Increased M5 labeling also corresponded with decreased unlabeled citrate and increased percent isotopomer labeling. Succinate, fumarate, and malate were mostly M2 and M4 labeled from ^13^C-glutamine labeling, as shown in Fig 5 for MDA-MB-23. Fig C in S1 Appendix shows the labeling for the other two cell lines.

**Fig 6.**
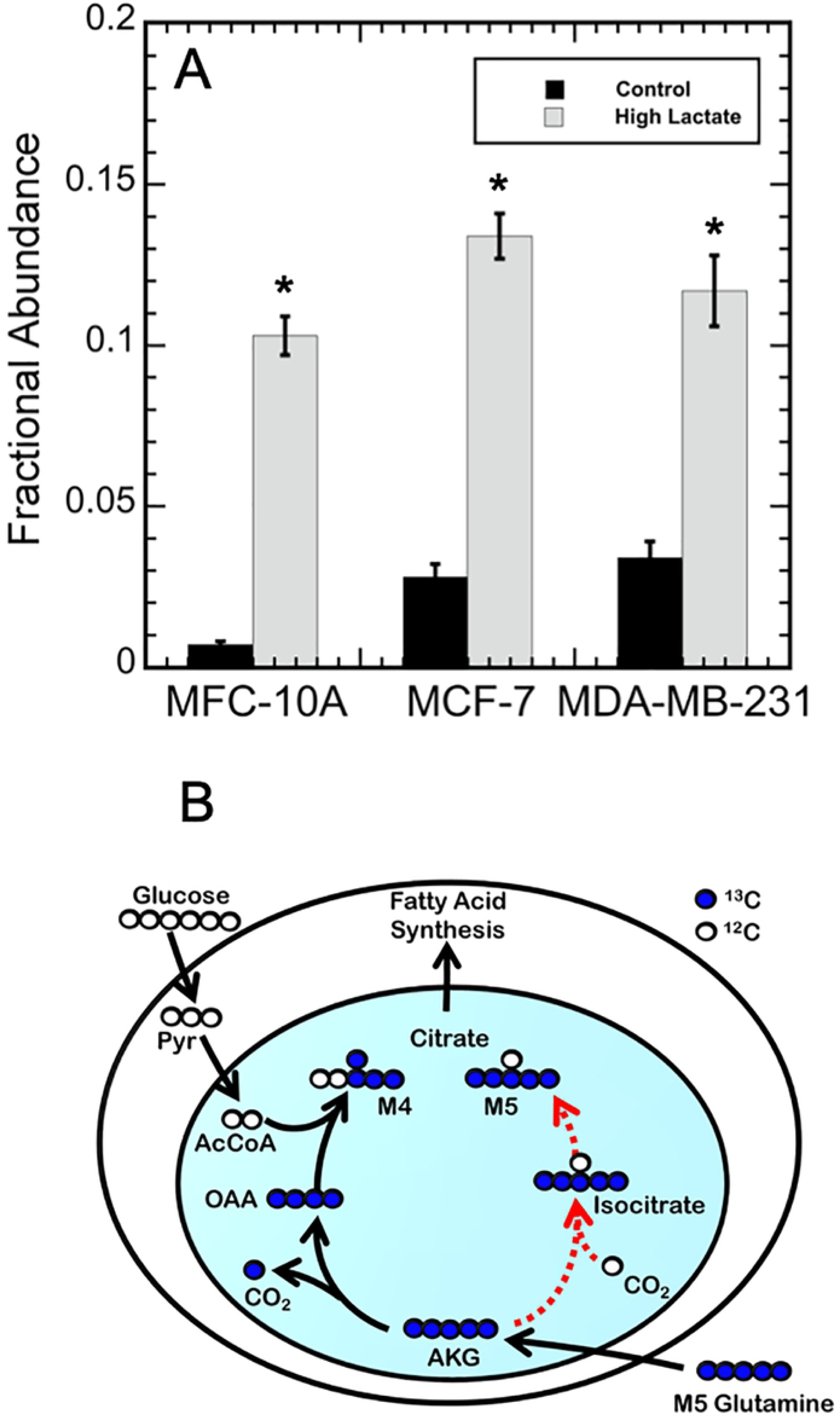
Evidence of reductive carboxylation of α-ketoglutarate (AKG) to citrate from [U-^13^C] glutamine labeling. A) Fractional abundance of M5 labeled citrate for MCF 10A, MCF7, and MDA-MB-231 cells from the control 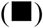 and high-lactate media 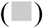. Error bars represent standard deviations. The astrick (*) indicates the fractional abundances were significantly different between the control and high-lactate media per cell line (p ≤ 0.01). B) Carbon atom transition for TCA metabolites from [U-^13^C] glutamine. The black lines depict oxidative glutamine metabolism that results in M4 citrate. The dashed red lines depict reductive glutamine metabolism that results in M5 citrate. For clarity, some reaction pathways were condensed.

### Metabolic flux analysis

To quantify the metabolic activity, Metran was used to predict the intracellular metabolite fluxes for each of the three cell lines under both conditions, six simulations in total. The experimental procedure section describes in detail the intracellular MIDs and extracellular fluxes used for the simulations. Each of the Metran model simulations converged to acceptable SSRes except for the MCF7 control condition, which slightly overfit. This overfitting was traced back to the high standard error for the glutamine MIDs from the biological replicates, where Tables E and F in S1 Appendix contain all the raw MID values for the three cell lines. This particular MID had a higher standard error than typically observed. Review of the data could not identify an experimental error, nor did this point meet outlier exclusion test criteria. Thus, this data point could not be discarded. Nevertheless, the effect of excluding this point on the model was examined to determine the effect this one data point had on the simulated flux values. When the Metran model simulation was conducted for the MCF7 control condition with only this metabolite MID standard error value replaced with the 0.6 mol% minimum machine error, the Metran simulation converged to an acceptable SSRes value. More importantly, there was no significant difference obtained for the simulated flux values. As the flux values are the critical outcome, the Metran simulations were conducted using the biological replicate MIDs (with the slight overfit SSRes) for the comparisons. The metabolic flux maps were generated for the three cell lines under each of the culture conditions, six in total using the actual biological replicates, as the flux outcomes were not significantly effected, eventhough the SSRes was overfit for MCF7. The flux maps are shown with standard deviations for the cell lines and conditions in Figs D-F in S1 Appendix. The list of metabolic fluxes for all Metran simulations are provided in Tables H-N in S1 Appendix, including the MCF7 simulation with the 0.6 mol% standard error for the single biological replicate.

As a part of the validation step, measured MIDs from the parallel glucose and glutamine tracer labeling experiments and extracellular flux measurements were compared to the simulated results from the model to further investigate the effects of lactate on metabolism and to determine cell line specific effects [21, 38]. As expected, glycolytic reactions exhibited the highest fluxes in both the control and high-lactate cultures. In all three cell lines, the glucose consumption and lactate production rates decreased due to high-lactate exposure. In addition, MFA simulations predicted slightly decreased PPP fluxes for all three cell lines due to high-lactate. As mentioned previously, increased M3 labeling of pyruvate and lactate from glutamine was observed for MCF 10A and MDA-MB-231 under high lactate, while MCF7 under high lactate resulted in a slight decrease in M3 labeling. This was reflected in the MFA simulations as increased malic enzyme fluxes, conversion of malate to pyruvate, for the MCF 10A and MDA-MB-231 cell lines under high lactate and a decreased malic enzyme flux for the MCF7 under high lactate.

## Discussion

In this study, the metabolic responses of three breast cell lines, MCF 10A, MCF7, and MDA-MB-231, to high-lactate were examined using ^13^C-MFA. MCF 10A is a non-tumorous epithelial breast cell line, MCF7 is a tumorigenic, luminal breast cancer cell line, and MDA-MB-231 is a metastatic, basal breast cancer cell line. ^13^C-glucose, ^13^C-glutamine, and ^13^C-lactate tracers were used in parallel experiments to determine the relative contribution of each substrate to intracellular metabolism. [1,2-^13^C] glucose and [U-^13^C] glutamine were selected as tracers, since both have previously been shown to be suitable for characterizing fluxes through glycolysis, the PPP, and TCA cycle pathways for mammalian cells [18, 21, 43, 44]. In addition, the [U-^13^C] lactate tracer was used to determine the metabolic contribution of lactate to breast cancer cell metabolism. For metabolic flux analysis, a detailed metabolic network model was constructed for breast cancer cells based a CHO cell metabolic network model [21]. The intracellular MIDs from parallel labeling experiments and extracellular flux measurements were paired with a mammalian metabolic network model to develop intracellular metabolic flux maps for each cell line under the control and high-lactate conditions.

As expected, the high-lactate condition caused a reduction in glucose consumption and lactate production for all cell lines examined (Fig 3). In addition, lactate consumption was observed in the high-lactate condition aided by the [U-^13^C] lactate tracer uptake in each cell line (Fig 4). Metabolic flexibility appears to play an important role in enabling cancer to thrive and survive in harsh environments, including inflammation, oxidative stress, and extreme nutrient and oxygen fluctuations [8, 45, 46]. As a result, the ability to metabolize higher levels of lactate offers potential advantages to cancer cells. The higher fractional abundance of intracellular M3 lactate labeling for MCF7 compared to MCF 10A and MDA-MB-231 in this study (Fig 4B) is consistent with previous finding that MCF7 display higher expression of monocarboxylate transporter 1 (MCT1), the lactate transporter associated with increased transport of lactate into the cell [6, 9]. Also, since TCA metabolites only had appreciable M1 labeling, this suggests that lactate was not a major source of acetyl-CoA and labeled metabolite pools were diluted by the metabolism of unlabeled pyruvate. Yet, the intracellular MIDs from the high-lactate cultures demonstrated that each cell line consumed and metabolized a significant amount of labeled lactate, which decreased both glucose consumption and net lactate production rates.

Intracellular metabolite labeling from [U-^13^C] glutamine was similar for the control and high lactate conditions for each cell line, with over 50% isotope labeling of all TCA cycle metabolites. Fig 5 shows the labeling for the MDA-MB-231 cells, where Fig C in S1 Appendix shows the labeling for all three cell lines. However, a slight increase in pyruvate and lactate labeling from glutamine was observed in MCF 10A and MDA-MB-231 due to high-lactate. This suggests that glutamine anaplerosis was enhanced to provide energy via oxidative phosphorylation, and then secreted as alanine and lactate in response to decreased glucose consumption and decreased energy generation via glycolysis. Interestingly, high-lactate also caused an increase in M5 citrate labeling for each cell line (Fig 6). The increased M5 citrate labeling suggests a rise in glutamine reductive carboxylation via the isocitrate dehydrogenase (IDH) reaction, as shown as a dashed red line in Fig 6B. Also, the decrease in M0 citrate suggests an increase in glutamine catabolism through the TCA cycle. Fig 5 shows the labeling for the MDA-MB-231 cells, where Fig C in S1 Appendix shows the labeling for all three cell lines. The MFA simulations also suggested cell line specific TCA cycle metabolic responses due to high-lactate, with TCA fluxes remaining unchanged for MCF 10A cells, slightly decreasing for MCF7 cells, and increasing for MDA-MB-231 cells. Furthermore, production of citrate from AKG via the IDH reaction – characterized by the exchange flux between AKG and citrate – increased for each cell line due to high-lactate. Altogether, these results suggest that the IDH reaction is highly reversible for each breast cell line tested.

Increases in fumarate, malate, and aspartate M3 labeling have also previously been observed with increased reductive carboxylation to support lipid synthesis in cancer cells grown under hypoxic conditions [18, 24, 25, 47-50]. This would result from [U-^13^C] glutamine serving as the main source of citrate via reductive carboxylation. Yet, M3 labeling of fumarate, malate, and aspartate did not significantly increase in this study. In this study, however, fumarate, malate, and aspartate were primarily M2 and M4 labeled, as shown in Fig 5 for the MDA-MB-231 cell line and shown in Fig C in S1 Appendix for all three cell lines. This indicates that the TCA metabolites were mainly derived from the glucose-derived pyruvate via oxidative metabolism rather than reductive glutamine metabolism. This implies that glutamine via reductive carboxylation was not the major carbon source for citrate synthesis in either condition. These observations agree with previous studies in which increased reductive carboxylation was observed for cancer cells without shifting the net IDH flux from oxidative to reductive glutamine metabolism in the TCA cycle [25, 49]. Jiang et al. 2016 also previously showed that increases in M5 citrate labeling without accompanied changes to M3 fumarate and malate labeling is indicative of reductive carboxylation for citrate and AKG shuttling rather than for lipid synthesis. By shuttling AKG and citrate from the mitochondria to the cytosol and back, cells can shuffle redox equivalents to maintain a net-oxidative TCA metabolism [25]. This might suggest that in high-lactate conditions, cancer cells utilize the reversible NADPH-depended IDH reactions (1 and 2) rather than the NAD+-dependent reaction (IDH3) to compensate for the loss in NAD+ generation as a result of decreased lactate production [25]. Recall, the two more aggressive breast cancer cell lines examined in this study maintained equivalent exponential growth rates to the control cultures at 20 mM lactate, while the MCF 10A cell line (a non-tumorigenic epithelial cell line) failed to grow in the 20 mM lactate condition, but could maintain equivalent exponential growth rates with the control cultures at 10 mM lactate.

Mammalian cells, including several cancer cell lines, have been observed to increase reductive carboxylation of glutamine under hypoxia [18, 47]. Reductive carboxylation has also been suggested to be a cellular response to environmental stresses, such as reactive oxygen species (ROS) regulation [25, 48]. In addition, Fendt et al. 2013 observed that supplementing media with 25 mM lactate resulted in decreased reductive carboxylation of glutamine for a lung carcinoma cell line (A549) grown in normoxia and a net consumption of lactate [48]; however, in the Fendt et al. 2013 studies, the growth rates of the cultures were not controlled or matched between the cultures exposed to lactate and those not exposed to lactate. In this study, the lactate concentrations were specifically selected such that the growth rates were the same between the control and high extracellular lactate cultures, i.e., the MCF 10A culture was exposed to 10 mM lactate, whereas the MCF7 and MDA-MB-231 were exposed to 20 mM lactate. Higher concentrations of lactate had been screened for all three cell lines, but equal growth rates were not maintained (data not shown). This major difference in the experimental set-up of cultures could explain the observation differences between this study and Fendt et al. 2013 with respect to lactate production. In this study, the lactate concentration was selected to be non-growth inhibitory, whereas in the previous studies, the effect of lactate on the growth rates is unknown. Therefore, this is the first observation of reductive carboxylation increasing due to high extracellular lactate in normoxia.

While cancer cells have long been known to produce high levels of lactate, especially in hypoxic tumor regions, cancer cells can also consume and metabolize lactate as a substrate [6, 9, 51]. The findings in this study highlight that high extracellular lactate induces a response mechanism in both non-cancer and cancer cell metabolism, which is similar to hypoxia or other mitochondrial stresses, even in the presence of sufficient oxygen. These results suggest that modulation of reductive carboxylation is a cellular metabolic response to stress, including high extracellular lactate and hypoxia. Additionally, the response observed is lactate dependent and cell line independent, under growth conditions where the cells can maintain exponential growth under the lactate stress.

## Acknowledgements

We would like to thank Dr. Brian Booth of Clemson’s Bioengineering Department for kindly provided to us with MDA-MB-231 cells. Additionally, we would like to thank Dr. David Bruce and Dr. Bethany Carter of Clemson’s Chemical Engineering Department for assistance with GC-MS analysis.

## Author contributions

**Conceptualization:** Arthur Nathan Brodsky, Sarah W. Harcum.

**Data curation:** Arthur Nathan Brodsky, Daniel C. Odenwelder, Sarah W. Harcum.

**Formal analysis:** Arthur Nathan Brodsky, Daniel C. Odenwelder, Sarah W. Harcum.

**Investigation:** Arthur Nathan Brodsky, Daniel C. Odenwelder.

**Methodology:** Arthur Nathan Brodsky, Daniel C. Odenwelder, Sarah W. Harcum.

**Resources:** Sarah W. Harcum.

**Visualization:** Arthur Nathan Brodsky, Daniel C. Odenwelder, Sarah W. Harcum.

**Writing – original draft:** Arthur Nathan Brodsky, Daniel C. Odenwelder, Sarah W. Harcum.

**Writing – review & editing:** Daniel C. Odenwelder, Sarah W. Harcum.

## Supporting information

S1 Appendix. Supporting information.

## References

1. Warburg O. On the origin of cancer cells. Science. 1956;123(3191):309–14. doi: 10.1126/science.123.3191.309. PubMed PMID: WOS:A1956ZQ16000002.

2. DeBerardinis RJ, Lum JJ, Hatzivassiliou G, Thompson CB. The biology of cancer: Metabolic reprogramming fuels cell growth and proliferation. Cell Metab. 2008;7(1):11–20. doi: 10.1016/j.cmet.2007.10.002. PubMed PMID: WOS:000252235100005.

3. Cairns RA, Harris IS, Mak TW. Regulation of cancer cell metabolism. Nat Rev Cancer. 2011;11(2):85–95. doi: 10.1038/nrc2981. PubMed PMID: WOS:000286506700009.

4. Heiden MGV, Cantley LC, Thompson CB. Understanding the Warburg Effect: The Metabolic Requirements of Cell Proliferation. Science. 2009;324(5930):1029–33. doi: 10.1126/science.1160809. PubMed PMID: WOS:000266246700031.

5. Kennedy KM, Dewhirst MW. Tumor metabolism of lactate: the influence and therapeutic potential for MCT and CD147 regulation. Future Oncol. 2010;6(1):127–48. doi: 10.2217/fon.09.145. PubMed PMID: WOS:000273635000017.

6. Kennedy KM, Scarbrough PM, Ribeiro A, Richardson R, Yuan H, Sonveaux P, et al. Catabolism of Exogenous Lactate Reveals It as a Legitimate Metabolic Substrate in Breast Cancer. PLoS One. 2013;8(9):20. doi: 10.1371/journal.pone.0075154. PubMed PMID: WOS:000326240100122.

7. Martinez-Outschoorn UE, Prisco M, Ertel A, Tsirigos A, Lin Z, Pavlides S, et al. Ketones and lactate increase cancer cell “stemness”, driving recurrence, metastasis and poor clinical outcome in breast cancer. Cell Cycle. 2011;10(8):1271–86. doi: 10.4161/cc.10.8.15330. PubMed PMID: WOS:000290145100023.

8. Pavlides S, Tsirigos A, Vera I, Flomenberg N, Frank PG, Casimiro MC, et al. Loss of stromal caveolin-1 leads to oxidative stress, mimics hypoxia and drives inflammation in the tumor microenvironment, conferring the “reverse Warburg effect” A transcriptional informatics analysis with validation. Cell Cycle. 2010;9(11):2201–19. doi: 10.4161/cc.9.11.11848. PubMed PMID: WOS:000279148400032.

9. Sonveaux P, Vegran F, Schroeder T, Wergin MC, Verrax J, Rabbani ZN, et al. Targeting lactate-fueled respiration selectively kills hypoxic tumor cells in mice. J Clin Invest. 2008;118(12):3930–42. doi: 10.1172/jci36843. PubMed PMID: WOS:000261237300017.

10. Bonuccelli G, Tsirigos A, Whitaker-Menezes D, Pavlides S, Pestell RG, Chiavarina B, et al. Ketones and lactate “fuel” tumor growth and metastasis Evidence that epithelial cancer cells use oxidative mitochondrial metabolism. Cell Cycle. 2010;9(17):3506–14. doi: 10.4161/cc.9.17.12731. PubMed PMID: WOS:000281621700024.

11. Stephanopoulos G. Metabolic Fluxes and Metabolic Engineering. Metab Eng. 1999;1(1):1–11. doi: 10.1006/mben.1998.0101. PubMed PMID: WOS:000208054500001.

12. Bonarius HPJ, Schmid G, Tramper J. Flux analysis of underdetermined metabolic networks: The quest for the missing constraints. Trends Biotechnol. 1997;15(8):308–14. doi: 10.1016/s0167- 7799(97)01067-6. PubMed PMID: WOS:A1997XP17900008.

13. Bonarius HP, Özemre A, Timmerarends B, Skrabal P, Tramper J, Schmid G, et al. Metabolic‐flux analysis of continuously cultured hybridoma cells using 13CO2 mass spectrometry in combination with ^13^C–lactate nuclear magnetic resonance spectroscopy and metabolite balancing. 2001;74(6):528–38.

14. Munger J, Bennett BD, Parikh A, Feng XJ, McArdle J, Rabitz HA, et al. Systems-level metabolic flux profiling identifies fatty acid synthesis as a target for antiviral therapy. Nat Biotechnol. 2008;26(10):1179–86. doi: 10.1038/nbt.1500. PubMed PMID: WOS:000259926000033.

15. Hiller K, Metallo CM. Profiling metabolic networks to study cancer metabolism. Curr Opin Biotechnol. 2013;24(1):60–8. doi: 10.1016/j.copbio.2012.11.001. PubMed PMID: WOS:000314487200009.

16. Nicolae A, Wahrheit J, Bahnemann J, Zeng AP, Heinzle E. Non-stationary ^13^C metabolic flux analysis of Chinese hamster ovary cells in batch culture using extracellular labeling highlights metabolic reversibility and compartmentation. BMC Syst Biol. 2014;8:15. doi: 10.1186/1752-0509-8-50. PubMed PMID: WOS:000335473900001.

17. Sauer U. Metabolic networks in motion: ^13^C-based flux analysis. Mol Syst Biol. 2006;2:10. doi: 10.1038/msb4100109. PubMed PMID: WOS:000243245400061.

18. Metallo CM, Walther JL, Stephanopoulos G. Evaluation of ^13^C isotopic tracers for metabolic flux analysis in mammalian cells. J Biotechnol. 2009;144(3):167–74. doi: 10.1016/j.jbiotec.2009.07.010. PubMed PMID: WOS:000272861600002.

19. Tang YJ, Martin HG, Myers S, Rodriguez S, Baidoo EEK, Keasling JD. Advances in analysis of microbial metabolic fluxes via ^13^C isotopic labeling. Mass Spectrom Rev. 2009;28(2):362–75. doi: 10.1002/mas.20191. PubMed PMID: WOS:000263580200007.

20. Templeton N, Dean J, Reddy P, Young JD. Peak antibody production is associated with increased oxidative metabolism in an industrially relevant fed–batch CHO cell culture. Biotechnol Bioeng. 2013;110(7):2013–24.

21. Ahn WS, Antoniewicz MRJMe. Parallel labeling experiments with [1, 2-13C] glucose and [U-13C] glutamine provide new insights into CHO cell metabolism. 2013;15:34–47.

22. Ahn WS, Antoniewicz MRJMe. Metabolic flux analysis of CHO cells at growth and non-growth phases using isotopic tracers and mass spectrometry. 2011;13(5):598–609.

23. Gaglio D, Metallo CM, Gameiro PA, Hiller K, Danna LS, Balestrieri C, et al. Oncogenic K-Ras decouples glucose and glutamine metabolism to support cancer cell growth. Mol Syst Biol. 2011;7:15. doi: 10.1038/msb.2011.56. PubMed PMID: WOS:000294537800005.

24. Metallo CM, Gameiro PA, Bell EL, Mattaini KR, Yang JJ, Hiller K, et al. Reductive glutamine metabolism by IDH1 mediates lipogenesis under hypoxia. Nature. 2012;481(7381):380–U166. doi: 10.1038/nature10602. PubMed PMID: WOS:000299210600046.

25. Jiang L, Shestov AA, Swain P, Yang CD, Parker SJ, Wang QA, et al. Reductive carboxylation supports redox homeostasis during anchorage-independent growth. Nature. 2016;532(7598):255-+. doi: 10.1038/nature17393. PubMed PMID: WOS:000374415100044.

26. Quek LE, Wittmann C, Nielsen LK, Kromer JO. OpenFLUX: efficient modelling software for C-13- based metabolic flux analysis. Microb Cell Fact. 2009;8:15. doi: 10.1186/1475-2859-8-25. PubMed PMID: WOS:000267250000001.

27. Weitzel M, Noh K, Dalman T, Niedenfuhr S, Stute B, Wiechert W. 13CFLUX2-high-performance software suite for 13C-metabolic flux analysis. Bioinformatics. 2013;29(1):143–5. doi: 10.1093/bioinformatics/bts646. PubMed PMID: WOS:000312654600027.

28. Young JD. INCA: a computational platform for isotopically non-stationary metabolic flux analysis. Bioinformatics. 2014;30(9):1333–5. doi: 10.1093/bioinformatics/btu015. PubMed PMID: WOS:000336095100033.

29. Zamboni N, Fendt SM, Ruhl M, Sauer U. 13C-based metabolic flux analysis. Nat Protoc. 2009;4(6):878–92. doi: 10.1038/nprot.2009.58. PubMed PMID: WOS:000266546200008.

30. Zancan P, Sola-Penna M, Furtado CM, Da Silva D. Differential expression of phosphofructokinase-1 isoforms correlates with the glycolytic efficiency of breast cancer cells. Mol Genet Metab. 2010;100(4):372–8. doi: 10.1016/j.ymgme.2010.04.006. PubMed PMID: WOS:000280440400011.

31. Fernandez CA, DesRosiers C, Previs SF, David F, Brunengraber H. Correction of 13C mass isotopomer distributions for natural stable isotope abundance. J Mass Spectrom. 1996;31(3):255–62. doi: 10.1002/(sici)1096-9888(199603)31:3<255::aid-jms290>3.0.co;2-3. PubMed PMID: WOS:A1996TZ91600004.

32. Yoo H, Antoniewicz MR, Stephanopoulos G, Kelleher JK. Quantifying reductive carboxylation flux of glutamine to lipid in a brown adipocyte cell line. J Biol Chem. 2008;283(30):20621–7. doi: 10.1074/jbc.M706494200. PubMed PMID: WOS:000257746100002.

33. Meadows AL, Kong B, Berdichevsky M, Roy S, Rosiva R, Blanch HW, et al. Metabolic and morphological differences between rapidly proliferating cancerous and normal breast epithelial cells. Biotechnol Prog. 2008;24(2):334–41. doi: 10.1021/bp070301d. PubMed PMID: WOS:000254703500007.

34. Bonarius HP, Hatzimanikatis V, Meesters KP, de Gooijer CD, Schmid G, Tramper JJB, et al. Metabolic flux analysis of hybridoma cells in different culture media using mass balances. 1996;50(3):299–318.

35. Azordegan N, Fraser V, Le K, Hillyer LM, Ma DW, Fischer G, et al. Carcinogenesis alters fatty acid profile in breast tissue. 2013;374(1-2):223–32.

36. Hilf R, Wittliff JL, Rector WD, Savlov ED, Hall TC, Orlando RA. Studies on certain cytoplasmic enzymes and specific estrogen receptors in human breast-cancer and in nonmalignant diseases of breast. Cancer Res. 1973;33(9):2054–62. PubMed PMID: WOS:A1973Q527800013.

37. Hilf R, Goldenberg H, Orlando RA, Archer FL. Some biochemical characteristics of human breast cancer and nonmalignant breast lesions. Proc Soc Exp Biol Med. 1969;132(2):613-+. PubMed PMID: WOS:A1969E661700048.

38. Leighty RW, Antoniewicz MR. Parallel labeling experiments with U-13C glucose validat. E. coli metabolic network model for 13C metabolic flux analysis. Metab Eng. 2012;14(5):533–41. doi: 10.1016/j.ymben.2012.06.003. PubMed PMID: WOS:000308369000009.

39. Antoniewicz MR, Kelleher JK, Stephanopoulos GJMe. Elementary metabolite units (EMU): a novel framework for modeling isotopic distributions. 2007;9(1):68–86.

40. Antoniewicz MR, Kelleher JK, Stephanopoulos GJMe. Determination of confidence intervals of metabolic fluxes estimated from stable isotope measurements. 2006;8(4):324–37.

41. Antoniewicz MR, Kelleher JK, Stephanopoulos GJAc. Accurate assessment of amino acid mass isotopomer distributions for metabolic flux analysis. 2007;79(19):7554–9.

42. Forbes NS, Meadows AL, Clark DS, Blanch HW. Estradiol stimulates the biosynthetic pathways of breast cancer cells: Detection by metabolic flux analysis. Metab Eng. 2006;8(6):639–52. doi: 10.1016/j.ymben.2006.06.005. PubMed PMID: WOS:000242137800011.

43. Crown SB, Ahn WS, Antoniewicz MR. Rational design of 13C-labeling experiments for metabolic flux analysis in mammalian cells. BMC Syst Biol. 2012;6:13. doi: 10.1186/1752-0509-6-43. PubMed PMID: WOS:000311224500001.

44. Crown SB, Antoniewicz MR. Selection of tracers for 13C-Metabolic Flux Analysis using Elementary Metabolite Units (EMU) basis vector methodology. Metab Eng. 2012;14(2):150–61. doi: 10.1016/j.ymben.2011.12.005. PubMed PMID: WOS:000301777400009.

45. Dayem AA, Choi H-Y, Kim J-H, Cho S-G. Role of oxidative stress in stem, cancer, and cancer stem cells. Cancers. 2010;2(2):859–84. doi: 10.3390/cancers2020859. PubMed PMID: MEDLINE:24281098.

46. Reuter S, Gupta SC, Chaturvedi MM, Aggarwal BB. Oxidative stress, inflammation, and cancer How are they linked? Free Radic Biol Med. 2010;49(11):1603–16. doi: 10.1016/j.freeradbiomed.2010.09.006. PubMed PMID: WOS:000284555000001.

47. Wise DR, Ward PS, Shay JES, Cross JR, Gruber JJ, Sachdeva UM, et al. Hypoxia promotes isocitrate dehydrogenase-dependent carboxylation of α-ketoglutarate to citrate to support cell growth and viability. Proc Natl Acad Sci U S A. 2011;108(49):19611–6. doi: 10.1073/pnas.1117773108. PubMed PMID: WOS:000297683800041.

48. Fendt SM, Bell EL, Keibler MA, Olenchock BA, Mayers JR, Wasylenko TM, et al. Reductive glutamine metabolism is a function of the α-ketoglutarate to citrate ratio in cells. Nat Commun. 2013;4:11. doi: 10.1038/ncomms3236. PubMed PMID: WOS:000323718300001.

49. Fan J, Kamphorst JJ, Rabinowitz JD, Shlomi T. Fatty Acid Labeling from Glutamine in Hypoxia Can Be Explained by Isotope Exchange without Net Reductive Isocitrate Dehydrogenase (IDH) Flux. J Biol Chem. 2013;288(43):31363–9. doi: 10.1074/jbc.M113.502740. PubMed PMID: WOS:000330778900053.

50. Grassian AR, Parker SJ, Davidson SM, Divakaruni AS, Green CR, Zhang XM, et al. IDH1 Mutations Alter Citric Acid Cycle Metabolism and Increase Dependence on Oxidative Mitochondrial Metabolism. Cancer Res. 2014;74(12):3317–31. doi: 10.1158/0008-5472.can-14-0772-t. PubMed PMID: WOS:000338877900012.

51. Pavlides S, Whitaker-Menezes D, Castello-Cros R, Flomenberg N, Witkiewicz AK, Frank PG, et al. The reverse Warburg effect Aerobic glycolysis in cancer associated fibroblasts and the tumor stroma. Cell Cycle. 2009;8(23):3984–4001. doi: 10.4161/cc.8.23.10238. PubMed PMID: WOS:000272219200038.

